# Transcranial Alternating Current Stimulation can disrupt or reestablish neural entrainment in a primate model of Parkinson’s disease

**DOI:** 10.1101/2025.09.17.676896

**Authors:** Harry Tran, Biswaranjan Mohanty, Noah Hjelle, Zhihe Zhao, Sangjun Lee, Adele DeNicola, Ivan Alekseichuk, Miles Wischnewski, Jing Wang, Jerrold Vitek, Luke Johnson, Alexander Opitz

**Affiliations:** Department of Biomedical Engineering, University of Minnesota, MN, USA; Department of Neurology, University of Minnesota, MN, USA; Stephen M. Stahl Center for Psychiatric Neuroscience, Department of Psychiatry and Behavioral Sciences, Feinberg School of Medicine, Northwestern University, IL, USA; Department of Experimental Psychology, University of Groningen, Netherlands

**Author notes:** <u>Corresponding authors</u> Harry Tran –, Alexander Opitz –. Co-first authors (Harry Tran, Biswaranjan Mohanty). Co-senior authors (Luke Johnson, Alexander Opitz).

**Keywords:** tACS, Parkinson’s disease, MPTP, single unit, phase locking

## Abstract

Transcranial alternating current stimulation (tACS) is a non-invasive brain stimulation method which can affect brain oscillations by inducing neuronal entrainment through modulation of spike timings. tACS has high potential for clinical applications in many neurological disorders in which the stimulation can be delivered to modulate oscillatory activity and disrupt pathological brain oscillations. For instance, in Parkinson’s Disease (PD), electrophysiological activity in the motor network often exhibits excessive and hyper-synchronized beta oscillations. However, the development of tACS as a therapeutic intervention for pathological oscillations requires the prior establishment of physiologically effective stimulation parameters. We recorded neuronal activity in the motor cortical area of three parkinsonian non-human primates and examined the influence of tACS-induced electric fields on neural firing patterns. We found that weak extracellular electric fields first disrupt beta-band spike timing patterns by changing the preferred spiking phase of neurons but eventually reestablish neural entrainment with altered phase preferences when electric fields are high. Additionally, we show that frequency-matched stimulation, when stimulation frequency corresponds to endogenous oscillatory activity, significantly enhances neural entrainment. Thus, tACS exhibits significant potential for controlling and modulating pathological oscillatory patterns in many neurological disorders such as PD.

## INTRODUCTION

Neurological disorders can exhibit pathological oscillatory patterns that interfere with healthy physiological brain rhythms, leading to impaired cognitive performance and disruption of neural information processing. In Parkinson’s disease (PD), these pathological oscillations are characterized by excessive synchronization of neuronal activity within the beta frequency band (13-30 Hz)^1–6^. Pathological oscillations can be found in both subcortical and cortical motor areas (premotor cortex PMC, primary motor cortex M1, and supplementary motor area SMA), and are thought to be related to impairment of motor functions, such as rigidity, bradykinesia, tremor, or posture instability^4,7,8^. Levodopa represents the primary medication-based therapy, though treatment efficacy varies among patients and disease stage. For patients with advanced PD, deep brain stimulation (DBS) targeting specific brain structures, such as the subthalamic nucleus (STN) or globus pallidus internus (GPi), has demonstrated significant therapeutic benefits in improving bradykinesia and reducing tremor severity^4,9,10^. These benefits are often accompanied by significant limitations, including surgical risks due to its invasive nature or potential hardware-related complications.

Non-invasive brain stimulation techniques are increasingly being explored as a treatment option for neurological disorders. Specifically transcranial alternating current stimulation (tACS) has gained attention from the scientific community due to its ability to modulate brain oscillations^11,12^. tACS applies oscillating currents between scalp electrodes to generate low-intensity electric fields that can modulate neural activity and induce synaptic plasticity^5,11–15^. tACS has been proposed as a treatment option for neurological disorders, including PD^16,17^, but its neural mechanisms remain incompletely understood, which hinder further translational developments. Animal and computational studies demonstrate that high-intensity tACS can induce neural entrainment - synchronization between neuronal spiking and the external stimulation oscillation^18–23^. While higher-intensity tACS effects have been independently validated^19,20^, lower-intensity tACS effects are the least understood despite being highly relevant for human applications^12,24^. One recent animal study showed that for low electric field strengths a competition for spike timing control occurs between endogenous neural oscillations and external stimulation^25^. This is further supported by a recent computational study which suggests that low-intensity tACS can desynchronize phase-locked neurons from endogenous oscillations^26^. Understanding these weak field effects is crucial for interacting with pathological brain oscillations such as the beta rhythm in PD. Depending on the applied electric field strength physiological effects will differ, affecting tACS outcomes. Elucidating these neuronal mechanisms would help optimize stimulation parameters such as intensity and montage, making tACS protocols more efficient to achieve a desired physiological response.

We performed multi-electrode electrophysiological recordings from cortical motor regions (M1, PMC, and SMA) in three non-human primates with toxin-induced parkinsonism to characterize the dose-response relationship of single-unit activity (SUA) during beta rhythm modulation (β-tACS). Stimulation was delivered at the prominent beta frequency (16 Hz) and at several control frequencies. The beta frequency was chosen to match the intrinsic oscillatory activity observed in the non-human primates used in this study. This investigation provides detailed insights into the neurophysiological mechanisms of tACS for disrupting pathological brain oscillations, such as beta activity in the case of Parkison’s disease.

## METHODS

### Animal preparation

All procedures were approved by the University of Minnesota Institutional Animal Care and Use Committee (IACUC) and complied with US Public Health Service policy on the humane care and use of laboratory animals. Three adult female rhesus macaques (Macaca mulatta, animal Ba, 17 years of age; animal Bu, 21 years; animal Pa, 19 years) were used in this study. Preoperative cranial Computed tomography (CT) and 7-T Magnetic Resonance Imaging (MRI) images were co-registered in the Monkey Cicerone neurosurgical navigation program^27^ and used for surgical planning for the placement of cortical arrays. Animal Ba was implanted with 96-channel Utah microelectrode arrays (Pt-Ir, 1.5 mm depth, 400 μm inter-electrode spacing, Blackrock Microsystems) using surgical methods described previously^28–30^. M1 and PMC were identified based on sulcal landmarks during the array implantation surgery, and an array was placed in each cortical target (Figure 1A, Supplementary Figure S1). Animal Bu and Pa were implanted with a 96-channel Gray Matter array loaded with tungsten microelectrodes with 1.5 mm spacing (Gray Matter Research, LLC), targeting the primary motor cortex (Figure 1A, right panel). Locations of the array and targeting areas were verified using fused pre-implantation MRI and post-implantation CT images.

**Figure 1.**
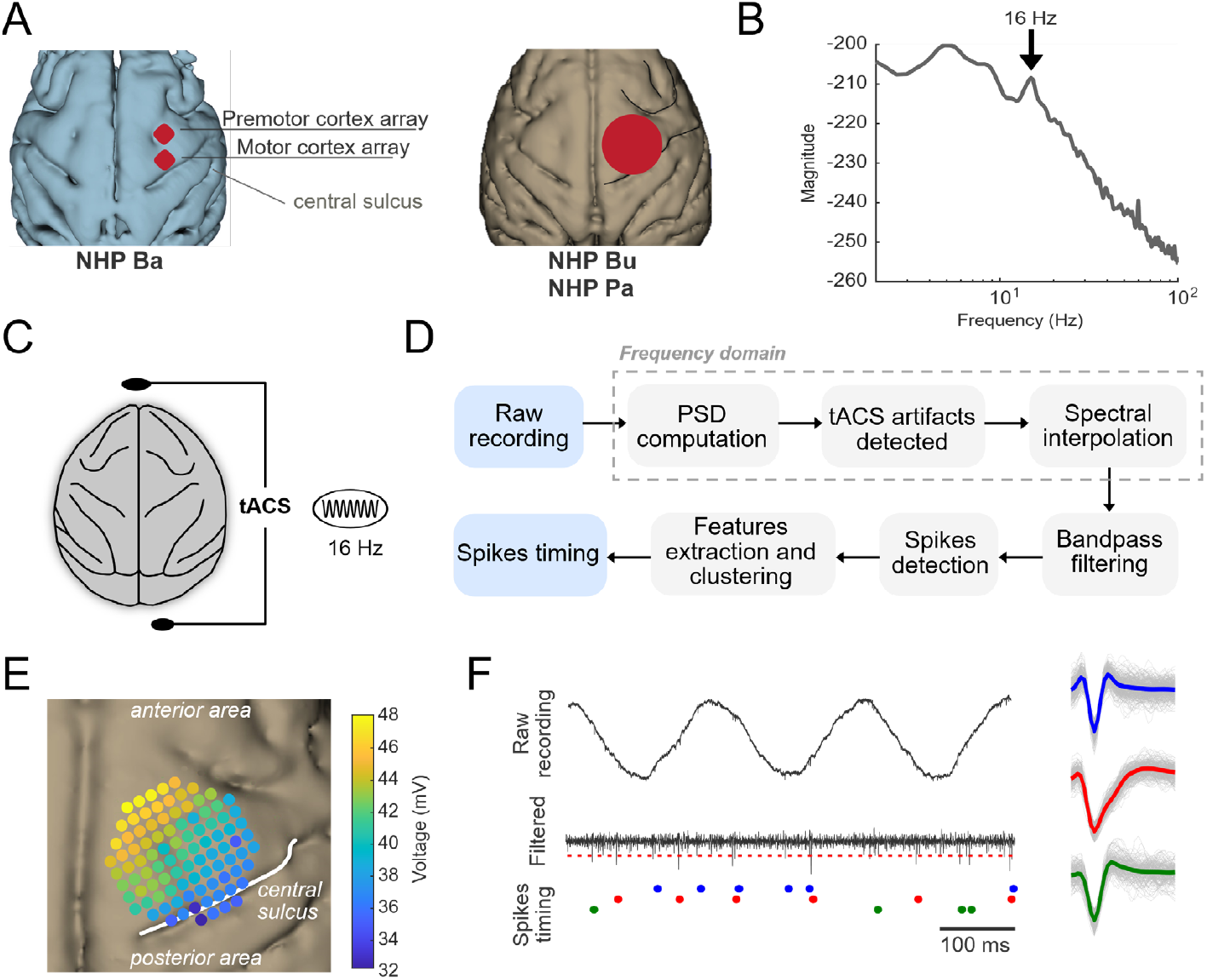
Overview of experimental design and data processing. (A) Two 96-channel Utah arrays (red square) were implanted over the premotor cortex (PMC) and motor cortex (M1) regions in animal Ba. A 96-channel Gray Matter Research grid array (red circle) was implanted in animals Bu and Pa over the motor areas (PMC, M1 and SMA). (B) Average of the power spectral density (PSD) of a recording session of Utah array channels where single units were detected in animal Ba. Parkinsonian state can induce a strong activity in cortical motor areas in the beta band (13-30 Hz). Black arrow highlights the endogenous brain oscillation around 16 Hz. (C) tACS was applied through two round electrodes (black ovals) attached to the frontal and occipital areas. The stimulation was thus applied in the anterior-posterior direction. (D) tACS artifacts were removed through PSD analysis of the raw data. Neural activity was subsequently identified by applying a threshold-based detection algorithm (red dashed line) to the artifact-free data after bandpass filtering between 300 and 3000 Hz. (E) Voltage gradient in animal Bu for 1 mA tACS. (F) Examples of three single units exhibiting various waveforms (triphasic in blue, monophasic in red and biphasic in green).

### MPTP Administration

Animals were rendered parkinsonian by intra-carotid and systemic intramuscular injections of the neurotoxin compound 1-methyl-4-phenyl-1,2,3,6-tetrahydropyridine (MPTP). Animal Ba received one intra-carotid and eight intramuscular injections (0.3-0.4 mg/kg each), Animal Bu received forty-two intramuscular injections (0.2-0.4 mg/kg each, a total of 1.8 mg/kg), and Animal Pa received twenty-seven intramuscular injections (0.2-0.4 mg/kg each). Overall parkinsonian severity was assessed using a modified Unified Parkinson’s Disease Rating Scale (mUPDRS), which involved the rating of axial motor symptoms (gait, posture, balance, turning, and defense reaction), food retrieval, and appendicular motor symptoms (upper and lower limb rigidity, bradykinesia, akinesia and tremor) on the hemi-body contralateral to neural recordings using a 0-3 scale (0 = normal, 3 = severe, maximum total score = 42) (Supplementary Table S1). The severity of the parkinsonian state was determined as mild, mUPDRS < 18; moderate, mUPDRS 18–31; and severe, mUPDRS ≥ 32^6^. For all subjects, data were gathered after a stable parkinsonian state was achieved.

### Neural activity recording

Neurophysiological data were collected using the Tucker-Davis Technologies (TDT, Alachua, FL, USA) workstation with a sampling frequency of ∼ 24 kHz. A large dynamic range (± 500 mV) and 28-bit resolution allowed recording of electrophysiological data during TACS, fully capturing both neural data and stimulation artifacts without data loss (e.g., due to amplifier saturation). Cortical neuronal activities were recorded while the animal was seated, head restrained in a primate chair without performing any type of task. No sensory stimulation of any kind was performed and the lights in the recording room were on. During each recording session, continuous behavioral monitoring was conducted to assess animal welfare.

### Electric stimulation

A sinusoidal electric current was applied through circular Pistim™ Ag/AgCl electrodes (contact area of *π* cm^2^) using the neurostimulator StarStim™ 8 system (Neuroelectrics®, Cambridge, MA, USA). There were two stimulation electrode arrangements (montages), inducing the anterior-posterior (AP) or left-right (LR) electric field direction. Stimulation electrodes were positioned at the frontal and occipital areas roughly corresponding to the Fpz and O1 coordinates in the human 10-20 EEG coordinates system for the antero-posterior (AP) direction (Figure 1C)^23^. For the left-right (LR) direction, electrodes were positioned on the anterior temporal areas just above the ears for each side, roughly corresponding to T7 and T8 in the human 10-20 system. A conductive gel (Ten20, Weaver and Company, CO, USA) was used, and impedances were checked before each stimulation session. We used a tACS stimulation frequency (*f*_*stim*_) of 16 Hz (β-tACS) corresponding to the endogenous frequency oscillation for the animals and a range of current intensities to investigate dose-dependent effects (0.05, 0.1, 0.2, 0.3, 0.4, 0.5, 0.75, and 1.0 mA). As control conditions, we employed two tACS frequencies (*f*_*stim*_): 5 Hz and 10 Hz, corresponding to theta and alpha bands, and applied currents to investigate dose-dependent effects (0.1, 0.2, 0.3, 0.5, 0.75, 1.0, and 1.5 mA). Overall, stimulation protocols involved 5 s or 10 s ramping up period (5 seconds for intensities below 0.5 mA and 10 seconds for intensities above 0.5 mA) followed by 4 minutes of stimulation period and then 5 or 10 s of ramping down period. Each recording session includes 4 minutes baseline recording (resting awake condition) followed by stimulation protocol with one minute of rest recording (no tACS stimulation) between two stimulation blocks.

### Electric field estimation

Electric field computation was performed through a multi-step signal processing approach. Raw electrode recordings were initially subjected to bandpass filtering using a second-order Butterworth filter centered on the applied stimulation frequency to isolate the tACS voltage signal (*f*_*stim*_ ± 1 Hz). The resulting voltage distribution across recording electrode locations was subsequently processed to estimate the electric field within the recording area. This estimation involved linear interpolation of recorded voltages onto a regular spatial grid, followed by numerical derivative across adjacent grid points to derive the electric field magnitude and direction (Figure 1E).

### Data preprocessing

The interaction between stimulation frequency *f*_*stim*_ and the neuromodulation device sampling rate (1000 Hz) induces a phenomenon called spectral leakage combined with aliasing effects. This phenomenon generates large predictable artifacts in the power spectral density (PSD) that require removal. These large predictable artifacts - observed at 1 kHz ± *f*_*stim*_ and their corresponding harmonics - were removed by applying a spectral interpolation ± 1 Hz around the artifact peak using FieldTrip MATLAB toolbox^31^ and then the PSD was computed again to make sure all peaks were removed. This preprocessing pipeline was applied until no high amplitude artifact peaks were observed. Once all artifacts’ peaks were removed, the recording channel was considered fully preprocessed and ready to be spike-sorted. The whole pipeline was performed on the Minnesota Supercomputing Institute’ (MSI) high performance computing of the University of Minnesota.

### Spike Sorting algorithm

We used the open-source MATLAB software Wave_Clus that is a semi-automatic algorithm for spike detection and sorting^32^. It uses wavelet decomposition to extract features of the spike waveforms and superparamagnetic clustering (SPC) to isolate the spikes in this feature space^32^. The raw recordings were first bandpass filtered using a 4^th^ order Butterworth filter between 300 and 3000 Hz and an amplitude thresholding based on the noise level was then applied to isolate single units.

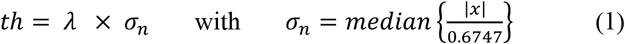

*x* is the bandpass-filtered signal, *σ*_*n*_ is an estimate of the standard deviation of the background noise^33^ and *λ* is a multiplying factor. Typically chosen values of *λ* are between 2 and 5, in our case, the value is 4^34–36^. Each spike was saved using 32 data points corresponding to 1.3 ms with a sampling rate of ∼ 24 kHz, 8 samples before the peak (negative or positive) and 24 after. All the spikes were saved in a matrix aligned with the peak.

The algorithm then computed the wavelet transform of the spike waveforms to extract features which are used as inputs for the clustering method (Figure 1F). For additional details, please refer to^32^.

### Quality of spike sorting and cluster isolations

*Inter-spike-interval (ISI)*. The ISI computes the time between subsequent spikes and can highlight specific trends in the firing behavior of a neuron (burstiness behavior for example). A thresholding in ISIs violation was used to distinguish single unit activity (SUA) and multi-unit activity (MUA). If 2 % of the ISIs were below 3 ms, the cluster was classified as a MUA; otherwise, it would be considered as a SUA^37^. In this study we only analyzed SUA.

*Waveform and variance*. It is assumed that each neuron fires spikes with a specific waveform that is stable over time. The standard deviation around the mean waveform should be stable in the 1.3 ms time window.

*Stability of amplitude over time*. The amplitude of the spike should remain stable over time and should not show any notable change during the different stimulation conditions.

*Signal-to-noise ratio (SNR)*. Finally, the SNR was calculated using the following formula^38^:

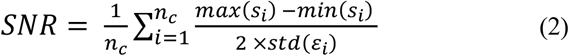

Where *s*_*i*_ is a vector of waveforms of spike *i, n*_*c*_ is the total number of spikes of cluster c, and 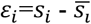 is the noise, 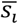 is the average spike waveform of cluster c.

### Neural entrainment quantification

We quantified neural entrainment by computing the phase-locking value (PLV) that estimates the phase of a spike timing relative to the phase of tACS. The PLV can be estimated by using the following formula^39^:

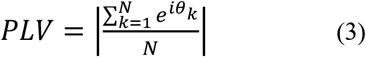

Where N is the number of action potentials and *θ*_*k*_ is the phase of the tACS stimulation at which the *k*_*th*_ action potential occurs. A PLV of 0 means that there is no synchronization while a value of 1 means perfect synchronization. This metric has been widely used and allows a direct comparison with other studies. During tACS, the local field potential (LFP) aligns with the shape of the stimulation, and the PLV is quantified using a filtered version of the LFP ± 1 Hz around the tACS stimulation frequency used using a 1st-order Butterworth filter. It is well established that a limited number of events can introduce bias in PLV by artificially increasing it so isolated single units with a low firing rate (< 0.5 Hz) were excluded from analysis^40^. For the baseline condition, PLV was computed with the filtered version of the LFP (*f*_*stim*_ ± 1 Hz).

Additionally, we use polar histograms to characterize the preferred spiking phase of neurons (0 degrees is the peak, 180 degrees is the trough of stimulation) for the different stimulation conditions. This approach quantifies the phase of tACS oscillation at the time a spike occurs. Under control conditions without external stimulation, neurons exhibiting stochastic firing patterns would be expected to demonstrate a uniform phase distribution. Data analysis was performed on a regular workstation (i7-9700K, 128 GB DDR4 3200 MHz, GPU RTX 2080) using custom MATLAB scripts.

### Statistical analysis

To test statistical significance of changes in PLVs and ΔPLVs, we implemented two generalized linear mixed models (glmm). For the changes in PLVs, the stimulation intensity (0.1, 0.2, 0.3, 0.5, 0.75 or 1 mA) and frequency (16 Hz AP, 10 Hz AP, or 5 Hz AP) or montage (16 Hz AP, 16 Hz LR) were the fixed effect factors, and the animal (Ba, Bu, or Pa) was the random effect factor. The distribution of PLVs was fitted to a gamma distribution and the model parameters were estimated with a log link function. Similarly, for the change in ΔPLVs, the electric field strength (V/m) and frequency or montage were the fixed effect factors, and the animal was a random effect factor. The distribution of ΔPLVs was fitted to a normal distribution and the model parameters were estimated with an identity link function. For the statistics, the F values, degree of freedom, and p-values are reported in the main text. We assessed whether individual neurons exhibited a significant preferred spiking phase using a Rayleigh test for non-uniformity using the circular statistics toolbox in MATLAB^41^. Significance in the change of firing rate across tACS intensities was evaluated by performing one-way ANOVA. The significance level was set at p = 0.05.

## RESULTS

We quantified the dose-response of β-tACS applied in the anterior-posterior direction on neural activity in three animals by calculating PLV under baseline and stimulation conditions (Figure 2A and 2B). A total of 88 SUA (Ba: 35, Bu: 22, Pa: 31 SUA) were successfully detected and isolated across the 16 Hz AP recording sessions. Among the 88 SUA recorded, approximately 56 % (n = 49) demonstrated significant phase-locking to the ongoing 16 Hz component of the baseline LFPs as determined by the Rayleigh test. Over the whole motor neuron population, the median PLV (reported here as median and 25-75% values) is 0.07 (0.04 - 0.10).

**Figure 2.**
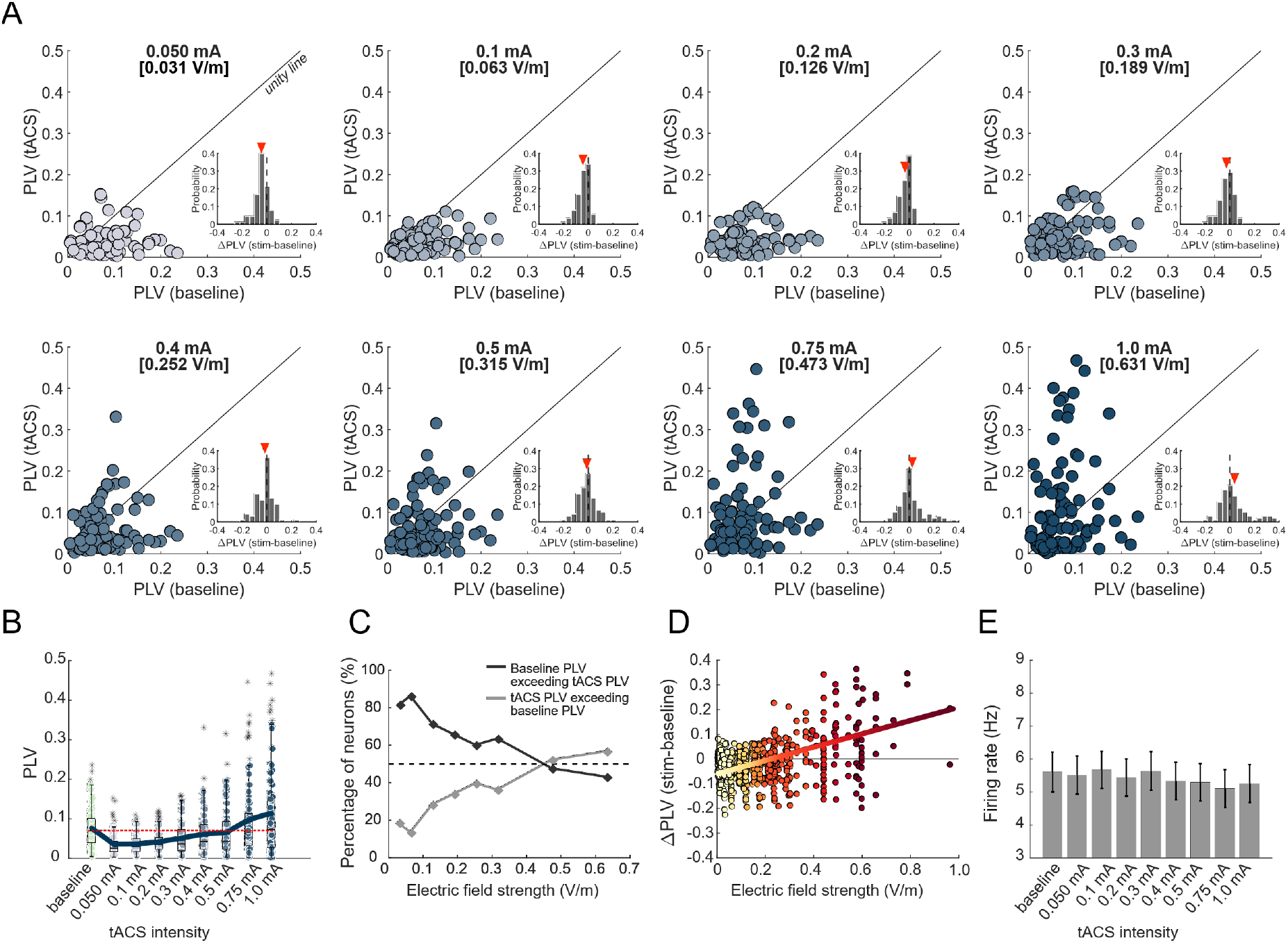
Effects of low electric field strength tACS in motor neurons. (A) Distribution of PLVs during 16 Hz tACS AP. Each dot compares the PLV of a single neuron for the baseline condition (x-axis) and the stimulation condition (y-axis). A dot above the unity line means that a neuron is more entrained during stimulation than baseline. Inset histograms show the difference between PLV at baseline and stimulation block ΔPLV - red arrow is the mean. For low intensity, the red arrow is negative, showing a lower neural synchronization compared to baseline. As tACS intensity increases the red arrow shifts towards a positive value, meaning a strong entrainment. (B) Boxplots of PLVs for the population across tACS intensities. We observe a linear increase of PLV with tACS intensity. Thick line is the mean (n = 88). Red dashed line is the median PLV at baseline. (C) Proportion of neurons that exhibit a higher (lower) PLV at baseline than stimulation condition in dark grey (light grey). (D) Distribution of ΔPLV with the electric field strength. (E) Firing rate has not shown any significant change over the baseline and stimulation condition (One-way ANOVA, F_7,696_ = 0.12, p = 0.9971). Vertical lines exhibit the standard error mean (SEM).

### Low-intensity β-tACS interferes with endogenous neural entrainment

We observed that weak electric fields modulate cortical neuron firing activity by disrupting spike timing synchronization within the beta frequency band (Figure 2A). This desynchronization phenomenon is mostly visible for tACS intensities below 0.2 V/m. Figure 2A compares the tACS PLV (y axis) with the baseline PLV (x axis) of each neuron (each dot) in a specific stimulation intensity. Dots below the unity line indicate neurons with higher PLV during baseline compared to tACS, while dots above the line indicate neurons with higher PLV during tACS compared to baseline. Among the 88 neurons, the majority exhibited a PLV at baseline that exceeded those observed during low intensity stimulation. Specifically, 81.6 % (n = 72), 86.4 % (n = 76), 71.6 % (n = 63), and 69.5 % (n = 58) of neurons demonstrated this pattern at electric field strengths of 0.03, 0.06, 0.13, and ∼ 0.2 V/m, respectively (tACS intensity of 0.05, 0.1, 0.2 and 0.3 mA) (Figure 2A). The unity line in Figure 2A represents the case where the PLV tACS is equal to the PLV baseline. The inset histograms in Figure 2A demonstrate the distribution of ΔPLV values (difference in PLV between stimulation and baseline conditions). At low electric field strengths (< 0.2 V/m), the histogram distribution is characterized by a negative shift, with the mean ΔPLV value (indicated by the red triangle) falling below zero. This disruption can also be observed for the same data set when the entrainment for low tACS intensities (≤ 0.3 mA) remains below the baseline condition (Figure 2B). Figure 2C shows the proportion of neurons with PLV above the unity line (dark grey) and below the unity line (light grey) as a function of stimulation intensity. PLV analysis revealed that a significant proportion of neurons maintained higher baseline values (dark grey line) below the electric field strengths of 0.3 V/m. Figure 2D shows that for weak electric field strengths (lower than ∼ 0.3 V/m), ΔPLV remains mainly negative, highlighting an electrophysiological phenomenon: a competition between endogenous and exogenous oscillations is happening for the control of spike timing. There was a minimal change in ΔPLV for the electric field below 0.3 V/m. These findings suggest that weak electric field strengths fail to induce neural entrainment, instead producing a perturbation that manifests as a decrease in PLV relative to the baseline condition. We investigated the change in firing rate with respect to stimulation intensities. One-way ANOVA revealed no significant effects of the β-tACS AP intensity on the firing rate (F_7,696_ = 0.12, p = 0.9971) (for individual tACS block, please refer to Supplementary Figure S2 and Supplementary Table 2).

### Dose-dependent effect of β-tACS

Neural entrainment gradually increased with tACS intensity (Figure 2A-D). A progressive increase in the proportion of neurons exhibiting statistically significant PLV was observed for tACS intensities above 0.4 mA. Specifically, significant PLV responses were demonstrated by 30.6 % (n = 27), 40.9 % (n = 36), and 46.6 % (n = 41) of neurons at electric field strengths of ∼ 0.3, 0.5, and 0.6 V/m, respectively. Additionally, inset histograms presented in Figure 2A demonstrate a systematic shift of the mean ΔPLV (denoted by the red triangle) toward positive values as a function of increasing electric field strength. This neural dose-response is additionally supported by the data presented in Figure 2B, where the mean PLV (blue curve) exhibits a monotonic increase as intensity increases. These findings are consistent with prior investigations demonstrating that electrophysiological effects can be induced at electric field strengths as low as 0.2 V/m^42,43^. We observed an increased proportion of neurons demonstrating enhanced PLV at higher electric field strengths (greater than 0.5 V/m, corresponding to tACS intensities above 0.8 mA). A minority of neurons (11.4 %, n = 10) exhibited PLV values greater than 0.3, while 3.4 % (n = 3) displayed PLV values exceeding 0.4 (Figure 2A, bottom panels). These findings are supported by the inset histograms (Figure 2A), which demonstrate that many neurons have a high ΔPLV above 0.2 (Figure 2A, bottom right panel; Figure 2D). Additionally, a linear regression was performed, and the slope was 0.27 (R^2^ = 0.57, p = 7.74 ×10^-19^) highlighting the dose dependency (Figure 2D).

### Low-intensity β-tACS alters the preferred phase of neuronal spike timing

At the baseline, recorded neurons exhibit phase-locking to the 16 Hz component of the LFP at a preferred spiking phase of 225.60° (Rayleigh test, p < 0.05) (Figure 3A). We analyzed the phase preferences of the neural population during tACS, with each data point representing an individual neuron (Figure 3B). This analysis linked the PLV and the preferred spiking phase of neurons. For an intensity below 0.3 mA, neurons exhibit minimal phase-locking: data points clustered near zero and the majority of neurons display PLV values below 0.07. For intensities above 0.3 mA, neurons exhibit a preferred spiking phase (∼ 180°) that becomes more prominent with high intensities.

**Figure 3.**
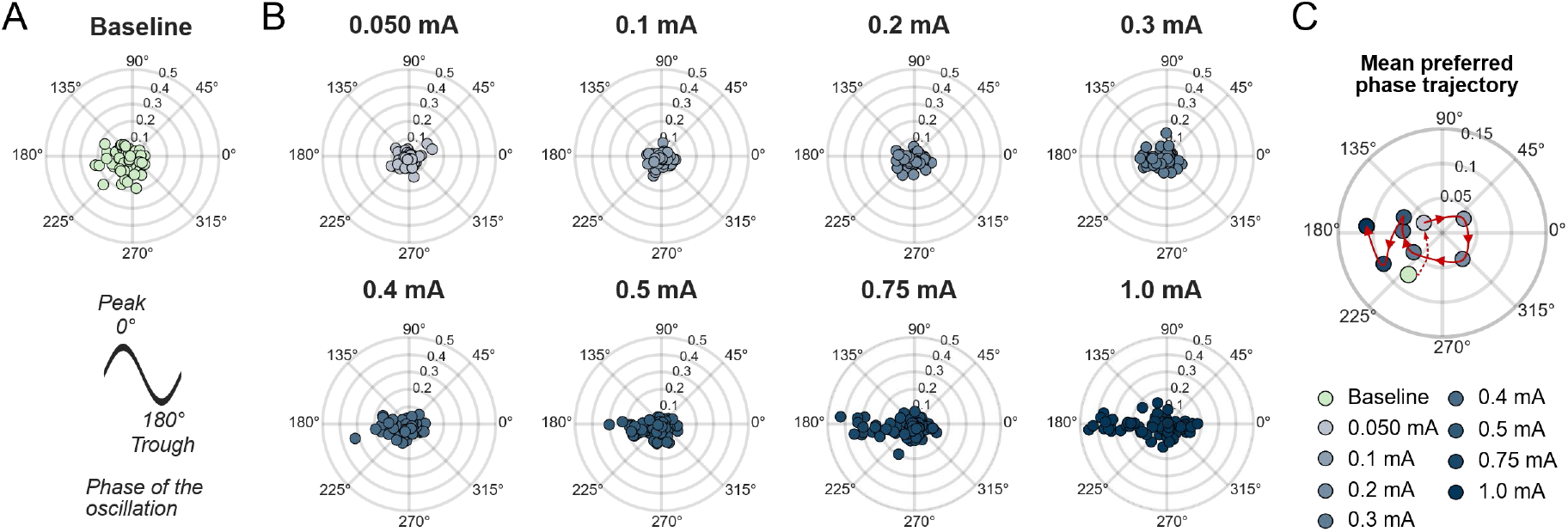
Low electric field strength tACS induces a shift in the preferred spiking phase. (A) Neuron populations exhibit a preferred spiking phase at baseline around 225° with a high PLV (approximately value is 0.07). The radius axis is PLV. The right panel shows the relation between the polar plot axis and the phase of the exogenous oscillations. (B) For low electric field strengths, the PLV is significantly reduced (dots are more concentrated and evenly distributed around 0). As tACS intensity increases, neurons tend to show a significant increase in PLV and exhibit a preferred spiking phase around 180° where the electric field strength is maximal. (C). Mean PLV and preferred spiking phase of the neuron population were computed to highlight the mean preferred phase trajectory. For low intensities, dots are close to 0 (highlighting the endogenous neural entrainment disruption). Neural entrainment is then reinstated with another preferred spiking phase when the electric field strength reaches a critical value.

We computed the mean spiking phase for each stimulation intensity, which is shown in Figure 3C and observed a preferred firing phase shift. The application of weak electric fields resulted in a substantial phase shift, with the mean phase decreasing from a baseline of 225.60° to 148.47° for a tACS intensity of 0.05 mA. This phase shift was accompanied by a corresponding decrease in the PLV, indicating reduced phase synchronization. The mean spiking phase demonstrates a large spread (Figure 3B and 3C) for low tACS intensities (0.05-0.3 mA), whereas the PLV remains relatively stable around 0.04. For stimulation intensities exceeding 0.4 mA, while the mean spiking phase oscillates around 180°, a concurrent increase in PLV was observed, reflecting enhanced neural synchronization (Figure 3B, bottom row). For examples of the preferred spiking phase of individual neurons, please refer to Supplementary Figure S3.

### Neurons exhibit selective responses to stimulation frequencies

To investigate the frequency-dependent tACS effects, we delivered 10 Hz tACS stimulation (alpha band) to two animals (Ba and Bu) and 5 Hz tACS stimulation (theta band) to two animals (Ba and Pa), both in the AP direction (Figure 4A). For the 10 Hz tACS condition, 47 neurons were successfully isolated (Ba: 21, Bu: 26 SUA). At baseline, the neurons exhibit a median PLV of 0.07 (0.04 - 0.10) (Figure 4B, left panel). Similarly, under 5 Hz tACS stimulation, 60 neurons were successfully isolated (Ba: 34, Pa: 26 SUA). At baseline, these neurons showed weak entrainment to the 5 Hz frequency component, with a PLV of 0.04 (0.02 - 0.06) (Figure 4C, left panel). At the baseline, neurons had a significantly higher entrainment in the alpha band than the theta band (Supplementary Figure S4).

**Figure 4.**
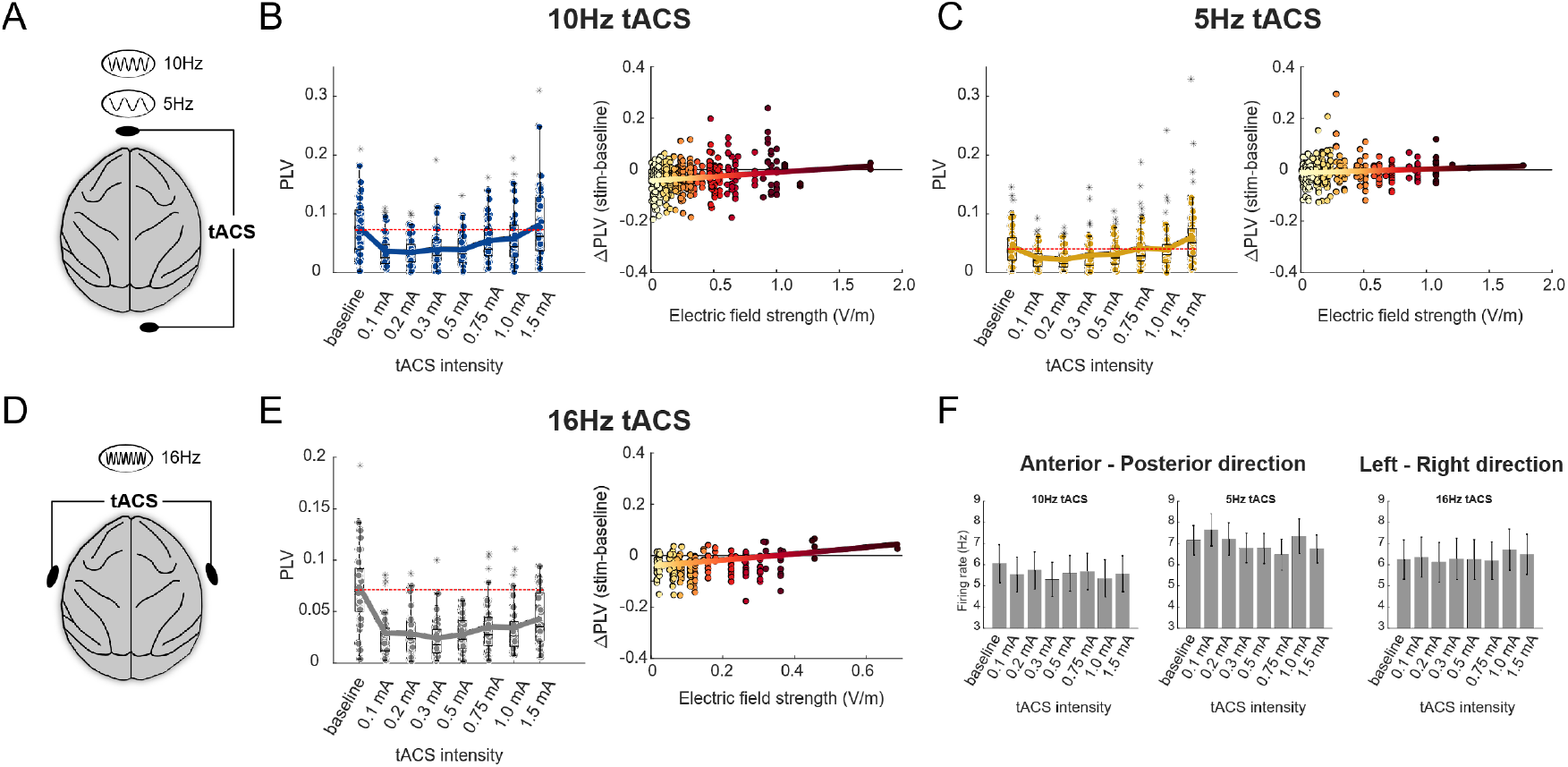
Neurons demonstrate frequency-selective and montage-specific responses. (A) Schematics of antero-posterior stimulation at frequencies of 10 Hz and 5 Hz. (B) *Left*: Boxplots of PLVs for the neuron population (n = 47) successfully isolated in two animals (Ba and Bu) across 10 Hz tACS intensities. *Right*: Distribution of ΔPLV with the electric field strength. (C) *Left*: Boxplots of PLVs for the neuron population (n = 60) successfully isolated in two animals (Ba and Pa) across 5 Hz tACS intensities. *Right*: Distribution of ΔPLV with the electric field strength. (D) Montage control condition includes a 16 Hz tACS in the left-right direction in animal Ba. (E) *Left*: boxplots of PLVs for the neuron population (n = 33) successfully isolated in animal Ba across 16 Hz tACS intensities. *Right*: Distribution of ΔPLV with the electric field strength. The red dashed line in the left panel of Figures B, C, and E represents the median PLV at baseline. The colored straight line in the right panels of Figures B, C, and E represents the linear regression fit. (F) Firing rates remain stable over the stimulation intensities regardless of the frequency or the montage stimulation. Vertical lines exhibit the standard error mean (SEM).

Across both 5 and 10 Hz control stimulation frequencies, we observed reduced PLVs at lower tACS intensity (< 0.4 mA) (Figure 4B-4C, left panel), suggesting that tACS intensity is not strong enough to induce significant entrainment. However, at higher intensities (above 0.5 mA), PLVs increased progressively with stimulation strength, revealing that tACS modulates spike timing in a dose-dependent manner (Figures 4B-4C, left panels). Additionally, we observed that in the higher tACS intensity (> 0.4 mA), the neural entrainment was stronger in β-tACS compared to control frequencies (10 Hz and 5 Hz). A comparative analysis of PLV distributions between 10 Hz tACS AP and β-tACS demonstrated that the change in PLVs was significant across frequency (glmm, F_1,621_ = 40.28, p = 4.25×10^-10^) and stimulation intensity (glmm, F_1,621_ = 126.8, p = 6.79×10^-27^). Similarly, the statistical analysis comparing 5 Hz tACS AP and β-tACS AP conditions shows that PLV changes were significant across both frequencies (glmm, F_1,753_ = 134, p = 8.95×10^-29^) and stimulation intensity (glmm, F_1,753_ = 103.99, p = 5.81×10^-23^).

Furthermore, we analyzed the relationship between electric field strength and ΔPLV (Figure 4B-4C, right panel). For the 10 Hz tACS, ΔPLV was mostly negative for low electric field strengths, highlighting neural desynchronization. The regression analysis depicted in Figure 4B (right panel) exhibited a lower slope coefficient compared to the corresponding analysis for β-tACS (Figure 2D), highlighting a frequency-specific entrainment effect (slope 0.27 vs of 0.03 for β-tACS and 10 Hz tACS, respectively). The change in ΔPLVs was significant across frequency (glmm, F_1,883_ = 6.15, p = 0.01) and electric field strength (glmm, F_1,883_ = 114.74, p = 2.95×10^-25^). For the 5 Hz tACS, ΔPLV was mostly negative and exhibited smaller magnitudes compared to the β-tACS or 10 Hz tACS. The smaller ΔPLV in 5 Hz tACS could be explained by the combination of two distinct neurophysiological phenomena: a low entrainment in the theta band at baseline, combined with a weak dose-dependency response resulting in lower ΔPLV (Figure 4C, right panel; slope of linear regression 0.01, R^2^ = 0.07, p = 0.0624). Statistical analysis revealed significant ΔPLV modulation compared to β-tACS (glmm, F_1,782_ = 28, p = 1.57×10^−7^) and electric field intensity (glmm, F_1,782_ = 85.21, p = 2.46×10^−19^).

### Neuronal entrainment is montage-specific

To understand how the electric field direction influences the modulation of neuronal firing patterns by the induced electric field, we recorded one animal (Ba) while stimulating at 16 Hz tACS in the left-right direction (LR) and compared it with the AP stimulation. We first analyzed the firing rate at baseline and investigated the dose-response of cortical neurons (n = 33) (Figure 4D and 4E). The population exhibited a baseline median PLV of 0.07 (0.05 - 0.09), and there were no significant changes compared to the baseline of the β-tACS AP recording.

For the LR montage, median PLV increased monotonically with tACS intensity however, the PLVs were below the baseline PLV, indicating disruption of neural synchronization for lower tACS intensities (Figure 4E, left panel). Additionally, higher PLVs were exhibited by AP montage compared to LR montage, indicating the neural entrainment was significantly more pronounced with the β-tACS AP condition, highlighting a montage-specific response (glmm, F_1,405_ = 240.19, p = 7.21×10^-43^). Furthermore, the change in PLVs was significant across the stimulation intensities (glmm, F_1,405_ = 76.56, p = 5.82×10^-17^).

Additionally, for both montages, we compared ΔPLV with respect to the electric field strength (Figure 2D and 4E, right panel). For LR montage, ΔPLV remains stable with negative values for electric field strengths lower than 0.3 V/m. When it reaches ∼ 0.3 V/m, ΔPLV exhibits positive values, highlighting the tACS induced entrainment (slope of the linear regression 0.12, R^2^ = 0.29, p = 0.001). The LR montage demonstrates a lower slope compared to the AP montage (slope: 0.27). The statistical analysis demonstrated significant changes in ΔPLVs across montage (glmm, F_1,459_ = 16.57, p = 5.52×10^-5^) and electric field strength (glmm, F_1,459_ = 153.44, p = 1.33×10^-30^).

### Firing rate does not change regardless of the stimulation parameters

We compared the firing rate across all tACS intensities during different stimulation conditions (frequencies and montages) and found no significant changes. We conducted one-way ANOVA to compare the effects of 10 Hz tACS and 5 Hz tACS intensity on firing rate (Figure 4F, left two panels). For both cases, the analysis revealed no significant effects of the tACS intensity on the firing rate (F_7,368_ = 0.07, p = 0.9993 for 10 Hz tACS and F_7,472_ = 0.28, p = 0.9606 for 5 Hz tACS) (Figure 4F, left panels). For individual tACS blocks, please refer to Supplementary Figures S7 and S8 for 10 Hz and 5 Hz tACS, respectively. Lastly, one-way ANOVA revealed no significant effects of the β-tACS LR intensity on the firing rate (F_7,256_ = 0.04, p = 0.9994) (Figure 4F, right panel). For individual tACS blocks, see Supplementary Figure S9. For the mean firing rate of the control experiments, see Supplementary Table 3, Supplementary Table 4 and Supplementary Table 5.

## DISCUSSION

In this study, we investigated the dose dependency of β-tACS in three parkinsonian non-human primates by performing in vivo recordings from motor cortical neurons. Our findings demonstrate: (1) weak extracellular electric fields modulate spike timing by disrupting neural entrainment with the endogenous oscillations and shifting the preferred phase of neuronal firing; (2) tACS at high intensities reinstates neural entrainment with a different preferred spiking phase; (3) an amplification network effect when the stimulation frequency matches the endogenous neural oscillation frequency and (4) β-tACS has stronger neural effects compared to other control frequencies. These findings advance our understanding of how tACS can be used to modulate neural oscillations in PD, and with potential applications extending to other neurological conditions characterized by pathological brain oscillations.

Initially it was expected that tACS would primarily act to enhance ongoing brain oscillations^11,44^. It is thought that with tACS, the time-varying electric field rhythmically de- and hyperpolarizes neural membranes, leading to changes in neural spike timing^5,22^. During the depolarization phase, action potentials occur at a higher likelihood than during the hyperpolarization phase^22^. In line with these ideas, we and others have demonstrated that with increased tACS intensity, alignment of spiking activity with respect to the external oscillation becomes more pronounced^20^. In the present study, however, we found that weak electric fields (< 0.3 V/m) reduce neural phase-locking compared to baseline in more than 70% of neurons. For an intermediate electric field (between 0.3 V/m and 0.4 V/m), this proportion decreases to 60 % of the neuronal population. For higher electric field strength (above 0.4 V/m), this proportion falls to ∼ 40 %. Our results add to emerging evidence that tACS at low intensities will first desynchronize the existing coupling between LFPs and spiking activity^25,26^. As simulation intensities increase, tACS will start to entrain spiking to the external oscillation. These findings suggest the possibility to develop tACS as a tool that can both increase and decrease ongoing spiking activity with respect to a specific oscillation. For example, these data highlight the ability of tACS to reduce neural entrainment to endogenous beta oscillations, neuronal activity that is often observed to be pathologically high in Parkinson’s disease. For weak electric field strengths, neither endogenous nor exogenous oscillations are strong enough to take control of the spike timings. Finally, when exogenous stimulation generates strong enough electric fields (greater than 0.4 V/m), neurons are entrained to the exogenous oscillation. Reduced entrainment also caused shifts in the preferred phase of neural firing at weak electric field strengths. In a strong electric field, neurons exhibited spiking activity synchronized to either the negative (trough) or positive (peak) phases of the exogenous oscillations. This suggests that altering electric field strength can modulate the preferred spiking phase of neurons, thereby controlling information flow within neuronal circuits^23^.

We tested 5 Hz and 10 Hz tACS alongside the main condition of 16 Hz, finding similar neural phenomena during low-intensity stimulation across all frequencies, consistent with previous animal studies where such disruption was not limited to stimulation frequency or a brain region^25^. However, the entrainment effects showed frequency-dependent differences, with cortical neurons displaying lower PLVs for 5 Hz and 10 Hz compared to 16 Hz at similar field strengths. This suggests that while low-intensity stimulation can disrupt neural activity across frequencies, the strength of entrainment may be enhanced at specific frequencies, possibly due to a resonance effect where β-tACS more effectively matches endogenous oscillation frequencies, enhancing network synchronization^42,45^. Neurons were more entrained to 10 Hz than 5 Hz stimulation, possibly because 10 Hz rhythms overlap with the low beta band in non-human primates^6^. Neuronal morphology and spatial organization influence neural responses to electric stimulation. Direct visualization of neuronal structures near recording electrodes remains challenging in vivo.

Neuronal responses showed a directional dependency for β-tACS, with significantly enhanced responsiveness during AP versus LR orientation. AP tACS induces an electric field that aligns with these preferred neuronal orientations, creating stronger depolarization and hyperpolarization in the neural membrane, potentially leading to modifications in cognitive function^46,47^. This specific response is explained by the underlying neural architecture: neurons are organized in columns that run perpendicular to the cortical surface, with their dendrites predominantly oriented in the AP direction. Additionally, the low PLVs even at substantial stimulation intensities for 16 Hz LR may be attributable to two potential mechanisms^43^. First, the spatial orientation of cortical neurons is not optimally aligned for responsiveness - cortical neurons demonstrate maximal sensitivity to electric fields oriented along the AP direction (see Figure 2A and 2B). Second, the heterogeneous tissue distribution of the animal within the cranium significantly influences current flow patterns. CT and T1/T2-weighted MRI revealed accumulations of muscle and adipose tissue in the lateral cranial regions (it is known that NHPs have a larger distribution of muscle on the sides compared to frontal and occipital areas) (Supplementary Figure 6). These tissues exhibit high electrical impedance, thus shunting stimulation current down and resulting in lower electric field strengths^24^ (Figure 4E, right panel).

Finally, we observed that the firing rate remains stable over the different stimulation conditions regardless of the stimulation parameters (frequency, intensity, or montage). This supports the idea that the neuronal responses observed are fundamentally governed by the phase-coupling dynamics between exogenous electrical stimulation and endogenous cortical oscillations and not a nonspecific increase in cortical excitability.

### Limitations and future work

This study mainly focused on low frequency tACS within three specific frequency ranges (delta, alpha, and beta bands), all maintained below 20 Hz. A recent human study investigated the effects of 1.0 mA tACS at different frequencies on bradykinesia, specifically comparing β-band stimulation (20 Hz) with γ-band stimulation (70 Hz) and demonstrated that β-tACS exacerbated bradykinetic symptoms. In contrast, γ-tACS modulates the plasticity of M1 in Parkinson’s disease^48^. The NHP research provides us with the flexibility to test multiple intensities and frequencies and find the optimal parameters to develop tACS-based therapies. In our research, only electrophysiology data were collected, limiting our ability to correlate the efficacy of tACS parameters (frequency and intensity) with motor behaviors. In future studies, we will investigate the optimal tACS parameters to improve pathological motor symptoms.

Previous research has demonstrated that tACS can induce synaptic plasticity^12,49,50^. In this investigation, stimulation blocks were short (< 4 minutes), necessitating further examination of whether extended stimulation durations could induce lasting effects on neural activity.

Another challenge in understanding tACS effects also depends on both the targeted brain region (e.g., motor cortex versus dorsolateral prefrontal cortex DLPFC) and the underlying brain state (healthy versus parkinsonian condition). The DLPFC, which plays a key role in executive function, demonstrates altered activity patterns in parkinsonian states compared to healthy conditions^51–56^. Due to differences in cytoarchitecture and connectivity, it is possible that DLPFC will respond differently to stimulation than motor cortex. Further DLPFC has different predominant oscillations, such as theta or alpha compared to M1. Thus, tACS parameters may need to be adapted to effectively engage DLPFC compared to M1. Investigating the effects of tACS across multiple brain regions would provide crucial insights into the therapeutic potential of tACS and would help establish whether the observed regional activity is specific to parkinsonian states.

Previous research has demonstrated that excitatory neurons exhibit greater sensitivity to electric field strength due to their larger morphological dimensions^22^, whereas inhibitory neurons demonstrate enhanced responsiveness within cortical network dynamics^26^. Future studies should investigate how different neuronal subtypes respond to β-tACS within the beta frequency band and whether distinct neural activity patterns can be identified.

Investigating the effects of weak electric fields on neural activity in deep brain structures within the basal ganglia-thalamocortical (BGTC) network, particularly the globus pallidus and subthalamic nucleus (STN), would be of critical importance. Indeed, pathological beta oscillations are much more present in that network and effects of β-tACS could be more significant^6^. Transcranial temporal interference stimulation (tTIS) offers a potential solution for stimulating these deep brain structures^57,58^.

In conclusion, our findings provide evidence supporting the hypothesis that weak oscillating electric fields modulate spike timing in motor neurons through a dual mechanism: initial neural entrainment disruption to the endogenous oscillation, accompanied by a shift in preferred spiking phase, followed by re-establishment of neural entrainment at a distinct phase for high electric field strengths. The control of brain oscillations through external stimulation promises substantial biomedical progress for addressing pathological oscillatory patterns such as excessive β-band activity observed in PD. While these results demonstrate the potential for tACS to modify neural synchronization patterns, further investigation of the neural mechanisms is necessary to establish its clinical long-term efficiency. Such research may prove instrumental in developing clinically tACS-based therapies to effectively alleviate motor symptoms and significantly improve the quality of life for patients with Parkinson’s disease.

## ACKNOWLEDGMENTS

The authors acknowledge the Minnesota Supercomputing Institute (MSI) at the University of Minnesota for providing resources that contributed to the research results reported within this paper. We additionally thank the team members who took care of the animals Claudia Hendrix, Hannah Baker, and Elizabeth McDuell as well as our veterinary and animal care colleagues at the University of Minnesota Research Animal Resources (RAR).

## AUTHOR CONTRIBUTION

HT: Conceptualization, Methodology, Data collection, Data analysis, Writing, Reviewing & editing BM: Animal handling, Data collection, Data analysis, Writing, Reviewing & editing

NH: Animal handling, Data collection, Data analysis, Reviewing & editing ZZ: Data analysis, Reviewing & editing

SL: Data analysis, Reviewing & editing AD: Animal handling

IA: Data collection, Reviewing & editing MW: Conceptualization, Reviewing & editing JW: Resources, Reviewing & editing

JV: Resources, Reviewing & editing, Funding Acquisition

LJ: Resources, Supervision, Writing, Reviewing & editing, Funding Acquisition

AO: Resources, Conceptualization, Supervision, Writing, Reviewing & editing, Funding Acquisition

