## Supplementary_materials for "Transcranial Alternating Current Stimulation can disrupt or reestablish neural entrainment in a primate model of Parkinson’s disease"

*Co-first authors

^†^Co-senior authors

Corresponding authors

Harry Tran – htran [at] umn.edu

Alexander Opitz – aopitz [at] umn.edu

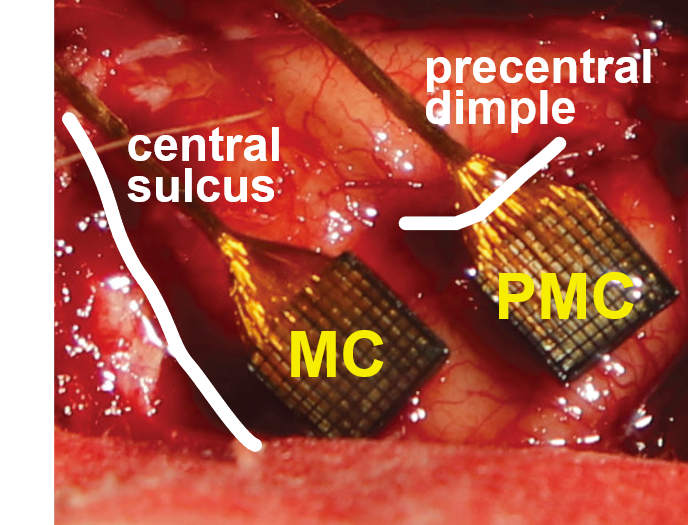

**Supplementary Figure 1. Location of the implantation of the Utah arrays.** Non-human primate Ba was implanted with two Utah arrays in the motor areas (motor cortex M1 and premotor cortex PMC). Anatomical characteristics such as sulcus are shown in white.

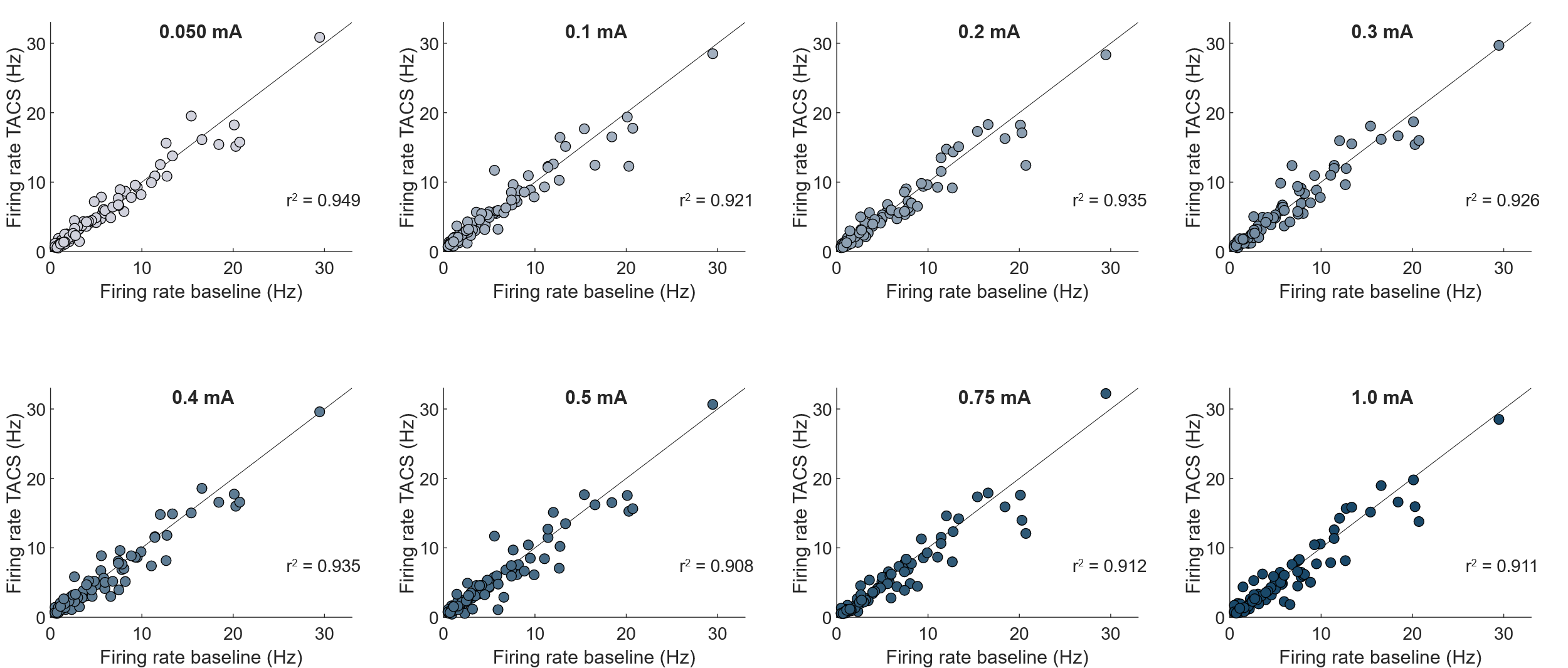

**Supplementary Figure S2. Firing rate does not show significant changes over TACS intensities for 16 Hz AP condition.** Neurons individual firing rate (represented by each individual dot) does not change between the baseline condition and the stimulation block – the closer a dot is to the unity line, the less change in firing there is. A linear regression was computed for each stimulation block and coefficient of correlation remains high (> 0.90).

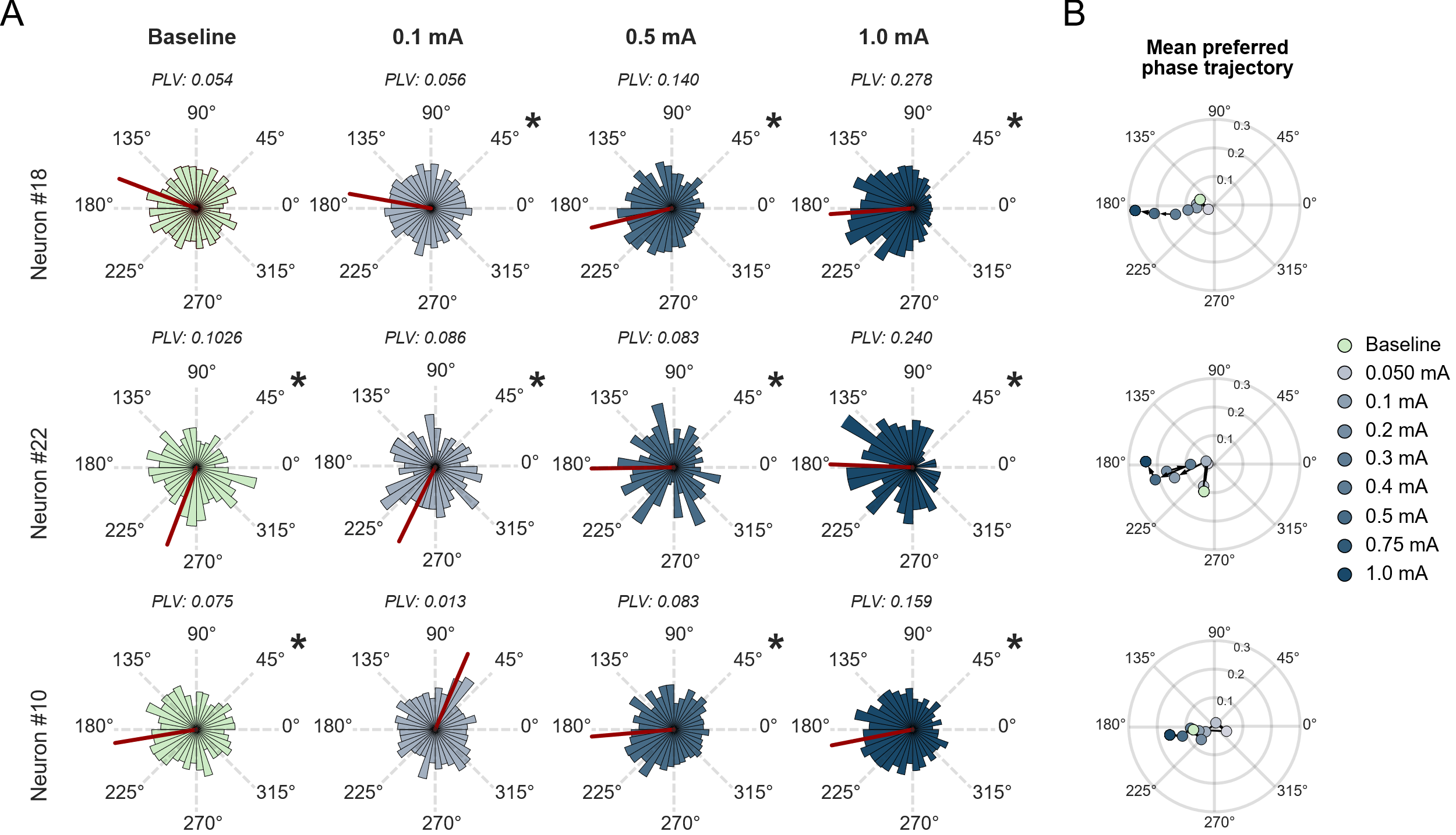

**Supplementary Figure 3. Example of behaviors of individual neurons for the 16 Hz stimulation AP.** (A) Neuron in the top row exhibits classic behavior: it becomes more entrained as TACS intensity increases. Neurons in the middle and bottom row exhibit neural desynchronization for an intensity of 0.1 mA TACS then become more entrained as TACS intensity increases. The red thick line represents the preferred spiking phase. A black star means significant entrainment (Rayleigh test, p < 0.05). (B) Mean preferred phase trajectory. Entrainment is reduced compared to baseline for low TACS intensities

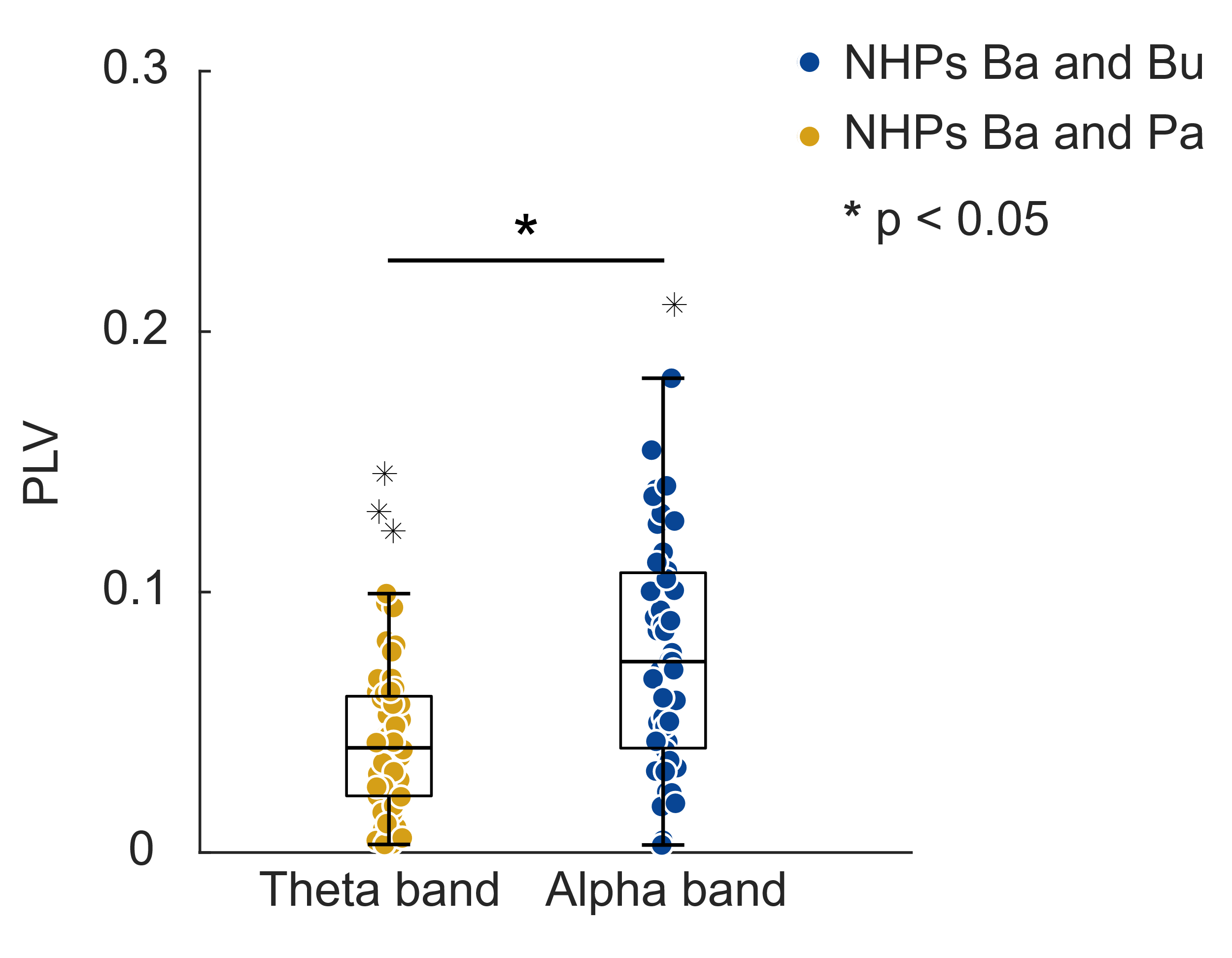

**Supplementary Figure 4. Neural entrainment at baseline in the theta and alpha band in three animals for control frequency condition.** Neurons in animals Ba and Bu are less entrained with the 5 Hz component of the LFP (n = 60) compared to those in animals Ba and Pa with the 10 Hz component of the LFP (n = 47). Black star indicates significant difference (two-sample *t*-test, t(105) = -4.3263, p = 3.464e-5).

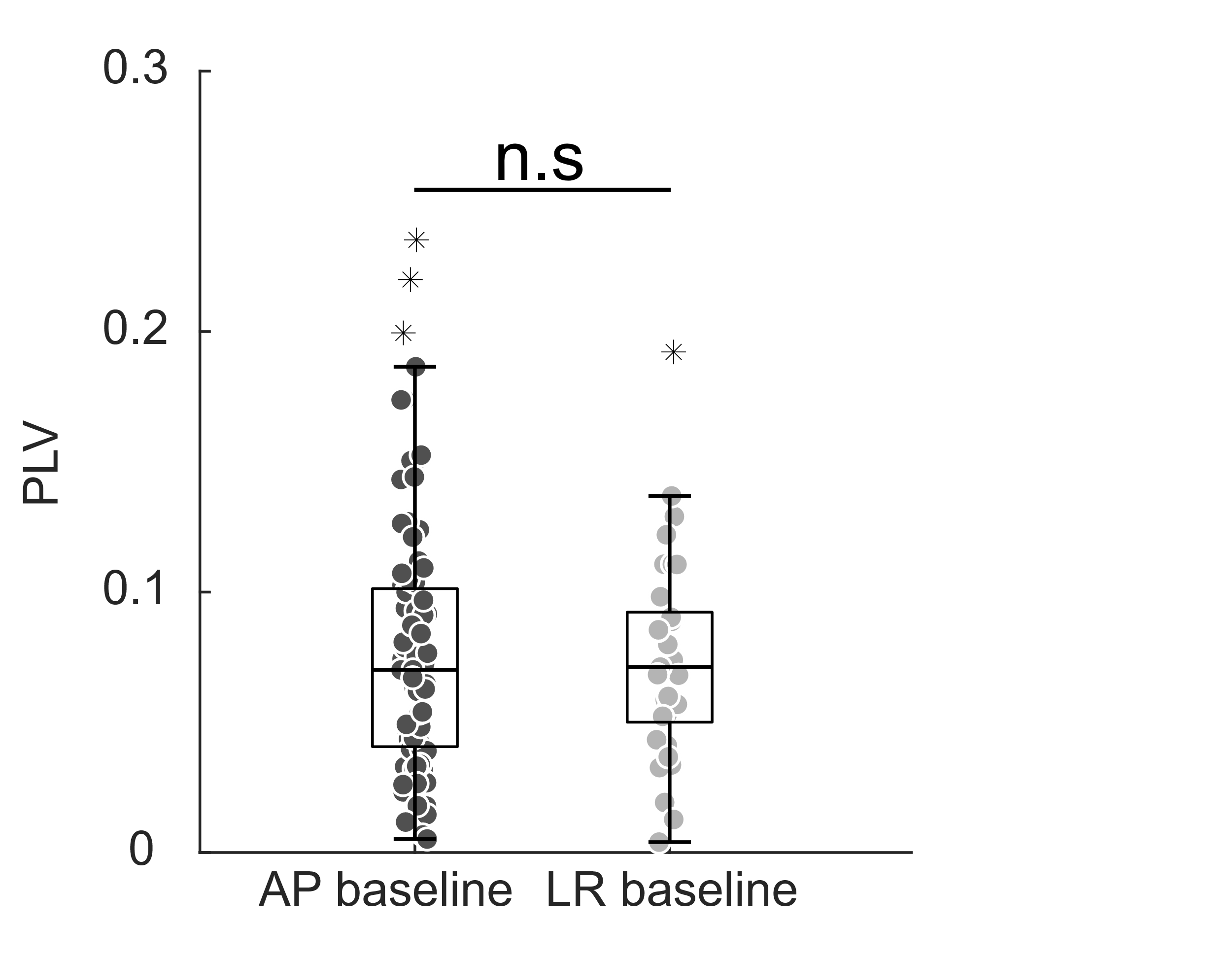

**Supplementary Figure 5. PLV of neurons in animal Ba for 16 Hz AP and 16 Hz LR for baseline condition.** The PLV of neurons at baseline did not show any significant differences between the two recording sessions of animal Ba. Results show no significant difference between the groups (t(119) = 0.3088, p = 0.7580). PLVs were 0.0769 ± 0.0488 for AP baseline (n = 88) and 0.0740 ± 0.0393 for LR baseline (n = 33) (mean ± standard deviation).

**Coronal view	Transverse view**

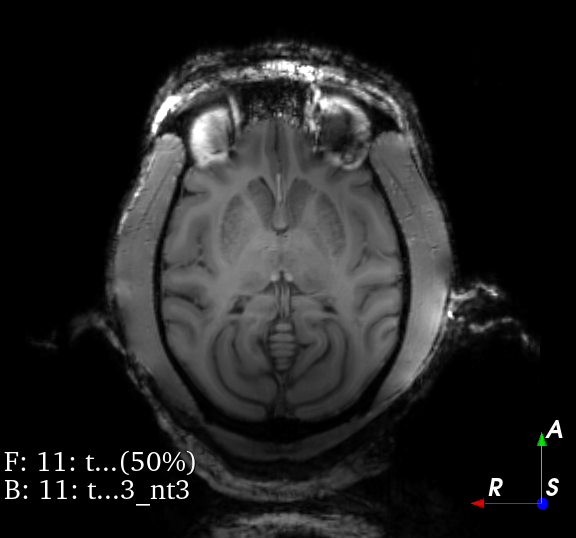

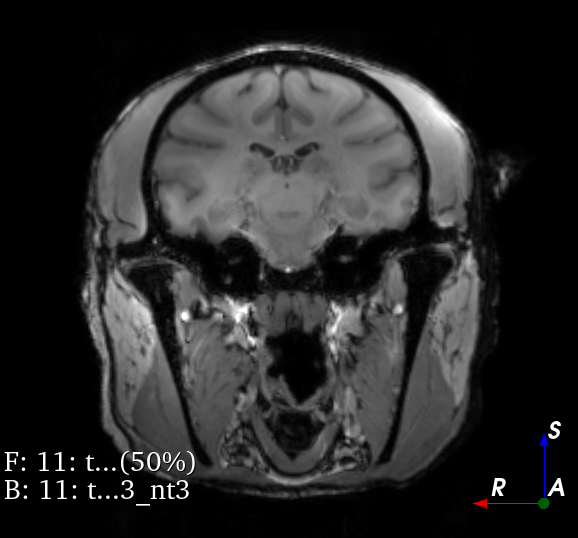

**Supplementary Figure 6. CT imaging of animal Ba.** A substantial bilateral distribution of muscle and adipose tissue is observed in the animal (pointed by white arrows), which accounts for the attenuation of the stimulation current.

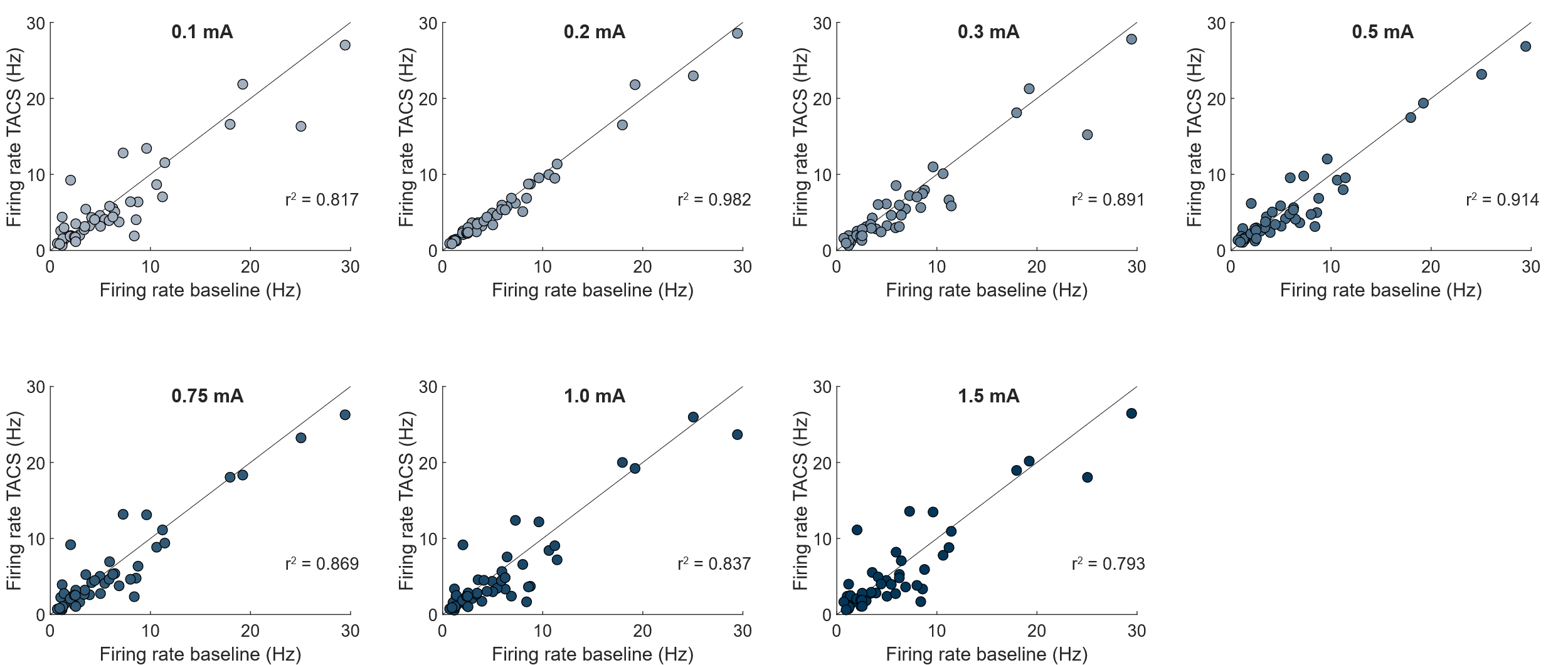

**Supplementary Figure S7. Firing rate does not show significant changes over TACS intensities for 10 Hz AP condition.** Neurons individual firing rate (represented by each individual dot) does not significantly change between the baseline condition and the stimulation block – the closer a dot is to the unity line, the less change in firing there is. A linear regression was computed for each stimulation block and coefficient of correlation remains high (> 0.70).

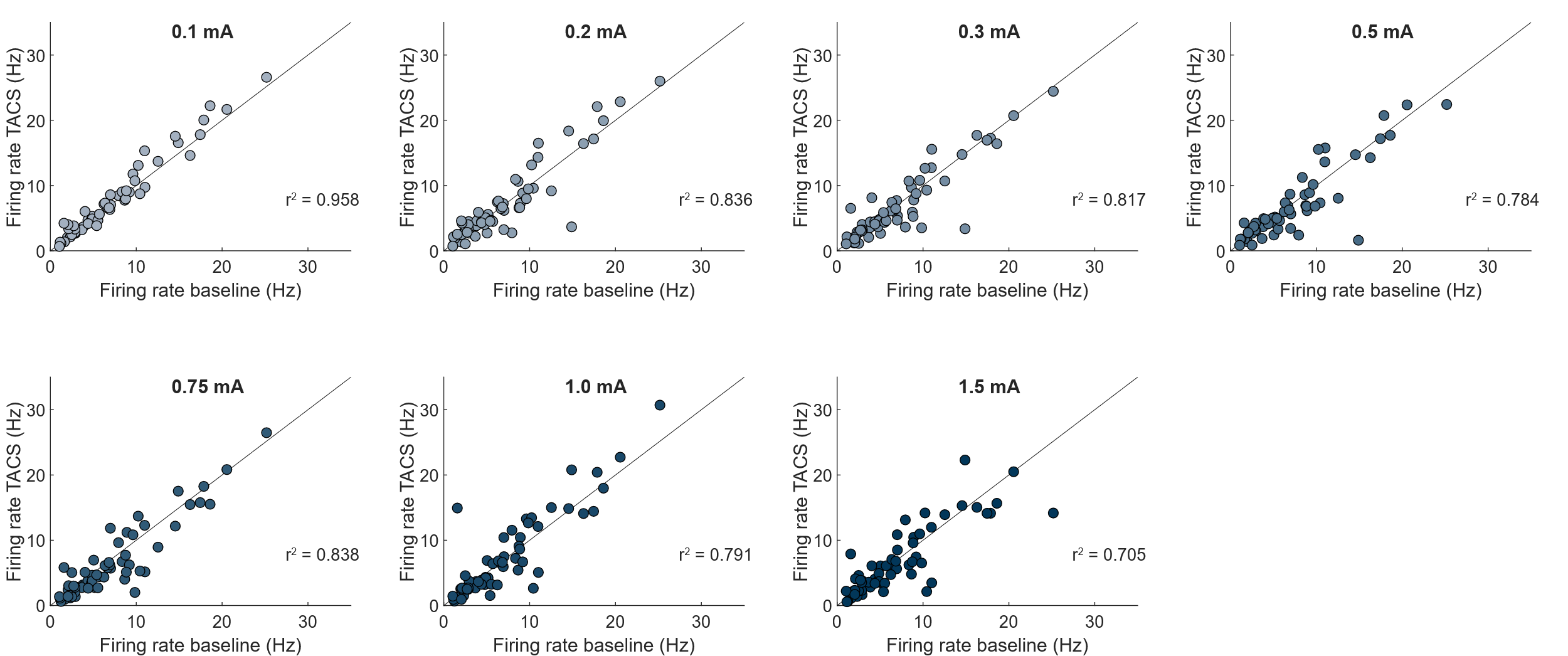

**Supplementary Figure S8. Firing rate does not show significant changes over TACS intensities for 5 Hz AP condition.** Neurons individual firing rate (represented by each individual dot) does not significantly change between the baseline condition and the stimulation block – the closer a dot is to the unity line, the less change in firing there is. A linear regression was computed for each stimulation block and coefficient of correlation remains high (> 0.70).

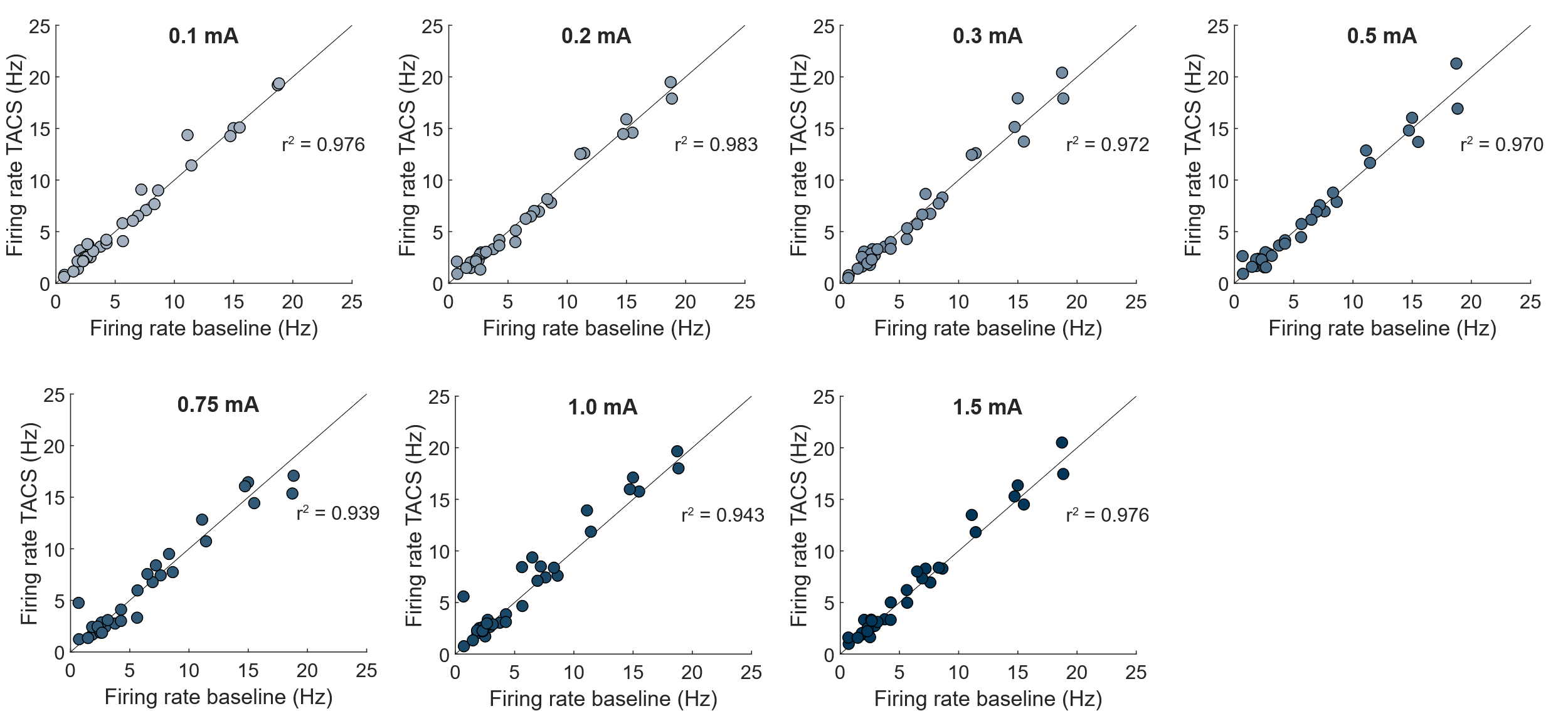

**Supplementary Figure S9. Firing rate does not show significant changes over TACS intensities for 16 Hz LR condition.** Neurons individual firing rate (represented by each individual dot) does not significantly change between the baseline condition and the stimulation block – the closer a dot is to the unity line, the less change in firing there is. A linear regression was computed for each stimulation block and coefficient of correlation remains high (> 0.90).

|  | Animal Ba | Animal Pa | Animal Bu |
| --- | --- | --- | --- |
| Motor Symptom |  |  |  |
| Rigidity (6) | 3 ± 0.4 | 4.5 ± 0.4 | 3.4 ± 0.1 |
| Tremor (6) | 1 ± 0.7 | 4.3 ± 0.9 | 2 ± 0 |
| Bradykinesia (6) | 4.1 ± 0.3 | 3.6 ± 0.5 | 3 ± 0 |
| Akinesia (6) | 2.5 ± 0.6 | 4 ± 0.4 | 3 ± 0 |
| Food Retrieval (3) | 2.1 ± 0.9 | 1.9 ± 0.5 | 1.5 ± 0 |
| Total (27) | 12.6 | 17.9 | 13.4 |
| Note: Number of sessions for Animal Ba (n = 4). Number of sessions for animal Pa (n = 2). Number of sessions for animal Bu (n = 2). | | | |

**Supplementary Table 1. mUDPRS for the three non-human primates.** Severity of symptoms was evaluated using a modified Unified Parkinson’s Disease Rating Scale (mUPDRS). Scores were computed on each recording day before applying the electrical stimulation. Animal Pa exhibit the most severe symptoms with an overall score of 17.9.

**Supplementary Table 2. Firing rate for the 16 Hz TACS AP conditions**

|  | TACS condition (mA) | | | | | | | |
| --- | --- | --- | --- | --- | --- | --- | --- | --- |
|  | 0.050 | 0.1 | 0.2 | 0.3 | 0.4 | 0.5 | 0.75 | 1.0 |
| Mean | 5.52 | 5.67 | 5.45 | 5.64 | 5.34 | 5.30 | 5.11 | 5.26 |
| Standard deviation | 5.35 | 5.24 | 5.26 | 5.44 | 5.27 | 5.30 | 5.36 | 5.33 |
| Note: Intensity does not have any significant effect on firing rate (one-ANOVA, p > 0.05) (n = 88) | | | | | | | | |

**Supplementary Table 3. Firing rate for the 10 Hz TACS AP conditions**

|  | TACS condition (mA) | | | | | | |
| --- | --- | --- | --- | --- | --- | --- | --- |
|  | 0.1 | 0.2 | 0.3 | 0.5 | 0.75 | 1.0 | 1.5 |
| Mean | 5.53 | 5.74 | 5.31 | 5.60 | 5.69 | 5.34 | 5.58 |
| Standard Deviation | 3.92 | 3.65 | 3.13 | 3.64 | 3.92 | 3.04 | 3.35 |
| Note: Intensity does not have any significant effect on firing rate (one-ANOVA, p > 0.05) (n = 47) | | | | | | | |

**Supplementary Table 4. Firing rate for the 5 Hz TACS AP conditions**

|  | TACS condition (mA) | | | | | | |
| --- | --- | --- | --- | --- | --- | --- | --- |
|  | 0.1 | 0.2 | 0.3 | 0.5 | 0.75 | 1.0 | 1.5 |
| Mean | 7.65 | 7.22 | 6.79 | 6.78 | 6.48 | 7.35 | 6.75 |
| Standard Deviation | 5.83 | 4.86 | 4.72 | 4.88 | 4.88 | 4.82 | 4.85 |
| Note: Intensity does not have any significant effect on firing rate (one-ANOVA, p > 0.05) (n = 60) | | | | | | | |

**Supplementary Table 5. Firing rate for the 16 Hz TACS LR conditions**

|  | TACS condition (mA) | | | | | | |
| --- | --- | --- | --- | --- | --- | --- | --- |
|  | 0.1 | 0.2 | 0.3 | 0.5 | 0.75 | 1.0 | 1.5 |
| Mean | 6.36 | 6.13 | 6.28 | 6.25 | 6.20 | 6.72 | 6.50 |
| Standard Deviation | 5.42 | 5.34 | 5.54 | 5.37 | 5.08 | 5.55 | 5.41 |
| Note: Intensity does not have any significant effect on firing rate (one-ANOVA, p > 0.05) (n = 33) | | | | | | | |
